# Impact of Chronic Total Occlusion Lesion Length onSix-month Angiographic and 2-year Clinical Outcomes

**DOI:** 10.1101/329268

**Authors:** Jihun Ahn, Seung-Woon Rha, ByoungGeol Choi, Se Yeon Choi, Jae Kyeong Byun, Ahmed Mashaly, Kareem Abdelshafi, Yoonjee Park, Won Young Jang, Woohyeun Kim, Jah Yeon Choi, EunJin Park, Jin Oh Na, Cheol Ung Choi Choi, EungJu Kim Kim, Chang Gyu Park, HongSeog Seo, DongJoo Oh, JinSu Byeon, SangHo Park, HyeYon Yu

## Abstract

**Background:** Successful chronic total occlusion (CTO) percutaneous coronary intervention (PCI) is known to be associated with improved clinical outcomes compared with failed CTO PCI. However, it is not clear whether the angiographic and clinical outcomes of long CTO lesionis different with those of short CTO lesion in the drug eluting stent (DES) era.

**Method sand Results:** A total of 235 consecutive patients underwent successful CTO intervention were divided into two groups according the CTO lesion length. Six-month angiographic and two-year clinical outcomes were compared between the two groups. The baseline clinical characteristics were similar between the two groups except prior PCI was more frequent in long CTO group whereas bifurcation lesion was more frequent in the short CTO group. In-hospital complications were similar between the two groups except intimal dissection was more frequent in long CTO group. Both groups had similar angiographic outcomes at 6 months and clinical outcomes up to 2 years except the incidence of repeat PCI, predominantly target vessel revascularization (TVR) was higher in long CTO group. In multivariate analysis, long CTO was an important predictor for repeat PCI (OR;4.26, CI 1.53-11.9, p=0.006).

**Conclusion:** The safety profile, angiographic and 2-year clinical outcomes were similar between the two groups except higher incidence of repeat PCI in long CTO group despite of successful PCI with DESs.

## Introduction

The use of drug-eluting stents (DES) and development of devices have result in reduction of restenosis and adverse events[1]. However, in spite of reduction of restenosis related with DES, clinical outcomes after percutaneous coronary intervention (PCI) is strongly affected by baseline lesion characters and long coronary artery lesion is a worse prognostic factor after PCI[2, 3].Also, despite the remarkable development of DES and remarkable progress of special devices and techniques in chronic total occlusion (CTO) PCI, procedural success rate of CTO patients is still lower than non-CTO patients and also the long term outcome is affected from clinical risk factors and baseline lesion characters[4]. However, there are no definite answers of CTO PCI outcomes according to the CTO lesion length, especially in DES era.

The aim of this study is to investigate whether the CTO lesion length can significantly impact on six-month angiographic and two-year clinical outcomes following successful CTO PCI.

## Methods

### Study population

A total of 235 consecutive CTO patients who underwent successful PCI with DESs from November 2004 to October 2010 at the Cardiovascular Center of Korea University Guro Hospital, Seoul, South Korea were enrolled for this study. Patients were divided into two groups according to the lesion length of the target vessel; The long CTO group (angiographic lesion length ≥30mm) and The short CTO group (angiographic lesion length<30mm).

This study protocol was approved by the institutional review board at Korea University Guro Hospital and all patients provided informed consent to participate in the study. The definition of CTO was a complete coronary obstruction with Thrombolysis In Myocardial Infarction (TIMI)flow grade 0 with an estimated duration ≥3 months with or without visible collateral flow[5]. The estimated duration of the CTO was determined by the interval from the last episode of angina symptoms consistent with the location of the occlusion. Only treated with de novo lesions were included and CTO due to in-stent restenosis (ISR) was excluded.

### Antiplatelet regimen

Loading doses of clopidogrel (300-600mg) and aspirin (200-300mg) were administered before the index procedure. Following PCI, all patients received aspirin (100mg) and clopidogrel (75mg) as maintenance dual antiplatelet regimen. Administration of clopidogrel was encouraged to continue at least for 1 year. Usage of adjunctive cilostazol to dual antiplatelet regimen (aspirin with clopidogrel) was depending on physicians’ discretion. Cilostazol was administered by 200mg post-loading and then 100mg twice a day for at least one month. Other concomitant medications were prescribed by physician’s decisions.

### PCI procedures

PCI procedures were performed with current guidelines using standard techniques. Various available guidewire were used to cross the CTO lesion and a variety of atheroablative devices were not utilized and mostly simple pre-dilation with balloon was performed to get an adequate luminal diameter which was necessary to accommodate the unexpanded stent and their delivery system. If necessary, adjunctive balloon dilatation was performed to achieve optimal outcome. All PCI was performed using DES and DES type was decided by the operator during the procedures. Procedural success was defined as a reduction in final angiographic diameter stenosis to less than 30% and TIMI grade≥2.

### Study end points and definition

#### Angiographic outcomes

Quantitative coronary angiographic (QCA) parameters were measured and analyzed before PCI, immediately after PCI and 6-months after the index procedure. The QCA measurement and analysis was performed at the Cardiovascular Center of Korea University Guro Hospital, Seoul. The primary angiographic end point was the incidence of binary restenosis at 6-12 months. Secondary angiographic end points were in-stent restenosis (ISR), mean diameter stenosis (DS), minimal luminal diameter (MLD) and late loss (LL) at 6 months after the index procedure.

### Clinical outcomes

Two-year after the index PCI, follow-up data was collected by face to face interview at out-patient clinic, review of medical record and/or telephone contact with patients. Primary clinical end point was the major adverse cardiovascular events (MACE)defined as the composite of total death, recurrent myocardial infarction (MI), and revascularization including PCI and coronary artery bypass graft (CABG). The secondary end points included death, MI, repeat PCI, target lesion revascularization (TLR), target vessel revascularization (TVR) and non-TVR. Deaths were considered as cardiac unless non-cardiac death could be confirmed. MACE was divided into TLR-MACE and TVR-MACE. TLR-MACE was defined as the composite of cardiac death, MI and TLR and TVR-MACE was defined as the composite of cardiac death, MI and TVR. Q-wave MI was defined as the development of new pathological Q-waves in at least two contiguous leads due to MI with or without an elevated creatinine kinase-MB fraction level.

### Statistical analysis

For continuous variables, differences between the two groups were evaluated by Student’s t-test. Data were expressed as mean ± standard deviations. Categorical variables were expressed as counts and percentages and analyzed with χ^2^ or Fisher’s exact test between groups as appropriate. The logistic regression model analysis was carried out to estimate for the risk of angiographic and clinical follow-up events with adjustment for risk factor such as sex, age, hypertension, diabetes mellitus (DM), chronic kidney disease, prior PCI and bifurcation lesion. A p-value of less than 0.05 was considered statistically significant. For all analyses, a 2-sided p < 0.05 was considered statistically significant. All statistical analyses were performed using SPSS 20 (IBM SPSS., Chicago, IL, USA).

## Results

### Baseline clinical, angiographic and procedural characteristics

In the present study, a total of 235CTO patients underwent PCI using DES were enrolled from November 2004 to October 2010.Theywere divided into two groups according to the lesion length of the CTO (long CTO group, ≥ 30mm: n=159, short CTO group, < 30mm: n=76). Baseline clinical and angiographic characteristics are demonstrated in Table 1.

**Table 1.** Baseline Clinical, Angiogrpahic and ProceduralCharacteristics

| Baseline characteristics, N (%) | Long CTO<br>(N=159) | Short CTO<br>(N=76) | P-value |
| --- | --- | --- | --- |
| **Clinical characteristics** |  |  |  |
| Male, sex | 119 (74.8) | 56 (73.6) | 0.848 |
| Age (years) | 60.94 ± 11.19 | 62.10 ± 10.78 | 0.446 |
| Left ventricular ejection fraction(%) | 49.24 ± 11.49 | 52.17 ± 11.18 | 0.076 |
| Prior myocardial infarction | 28 (17.6) | 5 (6.5) | 0.022 |
| Prior PCI | 40 (25.1) | 10 (13.1) | 0.035 |
| Prior CABG | 3 (1.8) | 1 (1.3) | 0.751 |
| Hypertension | 101 (63.5) | 46 (60.5) | 0.657 |
| Diabetes | 60 (37.7) | 26 (34.2) | 0.599 |
| Dyslipidemia | 49 (30.8) | 18 (23.6) | 0.257 |
| Cerebrovascular accident | 13 (8.1) | 5 (6.5) | 0.666 |
| Peripheral artery disease | 9 (5.6) | 2 (2.6) | 0.303 |
| Chronic Kidney disease | 11 (6.9) | 3 (3.9) | 0.368 |
| Smoker | 88 (55.3) | 38 (50) | 0.442 |
| **Angiographic and procedural characteristics** |  |  |  |
| Treated vessel per Patient | 1.949 ± 0.825 | 1.855 ± 0.795 | 0.401 |
| Treated CTO vessel per Patient | 1.075 ± 0.264 | 1.026 ± 0.161 | 0.137 |
| Multi-vessel disease | 101 (63.5) | 46 (60.5) | 0.657 |
| Multi-vessel CTO | 12 (7.5) | 2 (2.6) | 0.136 |
| CTO Lesion site |  |  |  |
| Left anterior descending artery | 74 (46.5) | 33 (43.4) | 0.653 |
| Left circumflex coronary artery | 31 (19.4) | 23 (30.2) | 0.066 |
| Right coronary artery | 66 (41.5) | 22 (28.9) | 0.062 |
| Stented CTO | 147 (92.4) | 73 (96) | 0.291 |
| LM disease | 8 (5) | 2 (2.6) | 0.393 |
| Bifurcation lesion | 37 (23.2) | 29 (38.1) | 0.017 |
| Total Lesion Length (mm) | 55.71 ± 27.16 | 21.56 ± 5.088 | < 0.001 |
| Vessel Size (mm) | 3.114 ± 0.438 | 2.920 ± 0.423 | <0.001 |
| Minimal lumen diameter(mm) | 2.726 ± 0.520 | 2.422 ± 0.679 | <0.001 |
| Acute gain (mm) | 2.726 ± 0.520 | 2.422 ± 0.679 | <0.001 |
| STENT diameter (mm) | 2.781 ± 0.361 | 2.861 ± 0.376 | 0.090 |
| STENT length (mm) | 29.83 ± 5.690 | 24.25 ± 5.626 | < 0.001 |
| Max Inflation (atm) | 14.40 ± 3.256 | 12.75 ± 3.178 | <0.001 |
| IVUS Guided PCI | 20 (12.5) | 12 (15.7) | 0.502 |
| Used stents on CTO lesion | 3.283 ± 0.948 | 2.986 ± 0.720 | < 0.05 |
| Used stent per Patient | 3.962 ± 1.391 | 3.644 ± 1.174 | 0.069 |
| Used stents (CTO lesion) |  |  |  |
| Sirolimus eluting stent (SES) | 43 (27) | 13 (17.1) | 0.094 |
| Paclitaxel eluting stent (PES) | 91 (57.2) | 22 (28.9) | < 0.001 |
| Zotarolimus eluting stent (ZES) | 30 (18.8) | 30 (39.4) | < 0.001 |
| Endeavor sprint | 15 (9.4) | 18 (23.6) | 0.003 |
| Endeavor resolute | 15 (9.4) | 12 (15.7) | 0.153 |
| Everolimus eluting stent (EES) | 14 (8.8) | 13 (17.1) | 0.062 |
| Promus &Xience | 13 (8.1) | 13 (17.1) | 0.041 |
| Promus Element | 3 (1.8) | 0 (0) | 0.228 |

CTO: chronic total occlusion; PCI: percutaneous coronary intervention; CABG; coronary artery bypass graft; iVUS:
Intravascular ultrasound.

The baseline clinical and angiographic characteristics were similar between the two groups except prior PCI(25.1% vs. 13.1%, p < 0.05) and prior MI (17.6% vs. 6.5%, p < 0.05)were more frequent in the long CTO group whereas bifurcation lesion (23.2% vs. 38.1%, p < 0.05) was more frequent in the short CTO group. Treated vessel size and stent diameter were smaller in the short CTO group.

### Clinical outcomes

Procedural and In-hospital complications were similar between the two groups except the incidence of significant intimal dissection was more frequent in the long CTO group, suggesting more complex and aggressive recanalization procedure (Table 2).

**Table 2.** Procedural Complications and In-Hospital Clinical Outcomes

|  | Long CTO<br>(N=159) | Short CTO<br>(N=76) | P-value |
| --- | --- | --- | --- |
| **Procedural complications, N (%)** |  |  |  |
| Perforation | 5 (3.1) | 0 (0) | 0.118 |
| Dissection | 34 (21.3) | 8 (10.5) | 0.042 |
| No reflow | 5 (3.1) | 1 (1.3) | 0.405 |
| Any hematoma(<4cm) | 1 (0.6) | 1 (1.3) | 0.591 |
| Major hematoma(>4cm) | 6 (3.7) | 5 (6.5) | 0.340 |
| Acute thrombosis | 0 (0) | 1 (1.3) | 0.147 |
| Acute renal failure | 1 (0.6) | 1 (1.3) | 0.591 |
| Cerebrovascular accident |  |  |  |
| **In-hospital clinical outcomes, N (%)** |  |  |  |
| Mortality | 2 (1.2) | 2 (2.6) | 0.446 |
| Cardiac death | 1 (0.6) | 2 (2.6) | 0.200 |
| Non-cardiac death | 1 (0.6) | 0 (0) | 0.488 |

Clinical outcomes for 2-year follow-up period are summarized in Table 3.

**Table 3.** Two-Year ClinicalOutcomes.

| Variables, N (%) | Long CTO<br>(N=159) | Short CTO<br>(N=76) | P-value |
| --- | --- | --- | --- |
| Mortality | 5 (3.1) | 4 (5.2) | 0.428 |
| Cardiac death | 2 (1.2) | 2 (2.6) | 0.446 |
| Non-cardiac death | 3 (1.8) | 2 (2.6) | 0.711 |
| Any Myocardial infarction | 3 (1.8) | 2 (2.6) | 0.711 |
| Q wave | 3 (1.8) | 2 (2.6) | 0.711 |
| Repeat PCI | 30 (18.8) | 6 (7.8) | 0.028 |
| TLR | 22 (13.8) | 5 (6.5) | 0.102 |
| TVR | 26 (16.3) | 5 (6.5) | 0.038 |
| Non TVR | 4 (2.5) | 0 (0) | 0.163 |
| All MACE | 34 (21.3) | 10 (13.1) | 0.130 |
| TLR MACE | 23 (14.4) | 7 (9.2) | 0.258 |
| TVR MACE | 30 (18.8) | 10 (13.1) | 0.275 |

PCI: percutaneous coronary intervention; TLR: target lesion revascularization; TVR: target vessel revascularization; MACE; major adverse cardiac event.

The incidence of adverse clinical events including total death (3.1%, 5.2%, p = 0.43), cardiac death (1.2% vs. 2.6%, p = 0.45), MI (1.8% vs. 2.6%, p = 0.71) and MACE (21.3% vs. 13.1%, p = 0.13) was not different between the two groups. However, the incidence of repeat PCI (18.8% vs. 7.8%, p < 0.05), predominantly TVR (16.3% vs. 6.5%, p < 0.05) was higher in the long CTO group.

The logistic regression model analysis was performed to estimate the risk of repeat PCI and TVR. Table 4 showed the predictors for repeat PCI and TVR. Long CTO was an important independent predictor for repeat PCI (OR; 4.26, CI 1.53 - 11.9, p <0.05) and TVR (OR; 4.23, CI 1.52 – 11.8, p < 0.05).

**Table 4.**
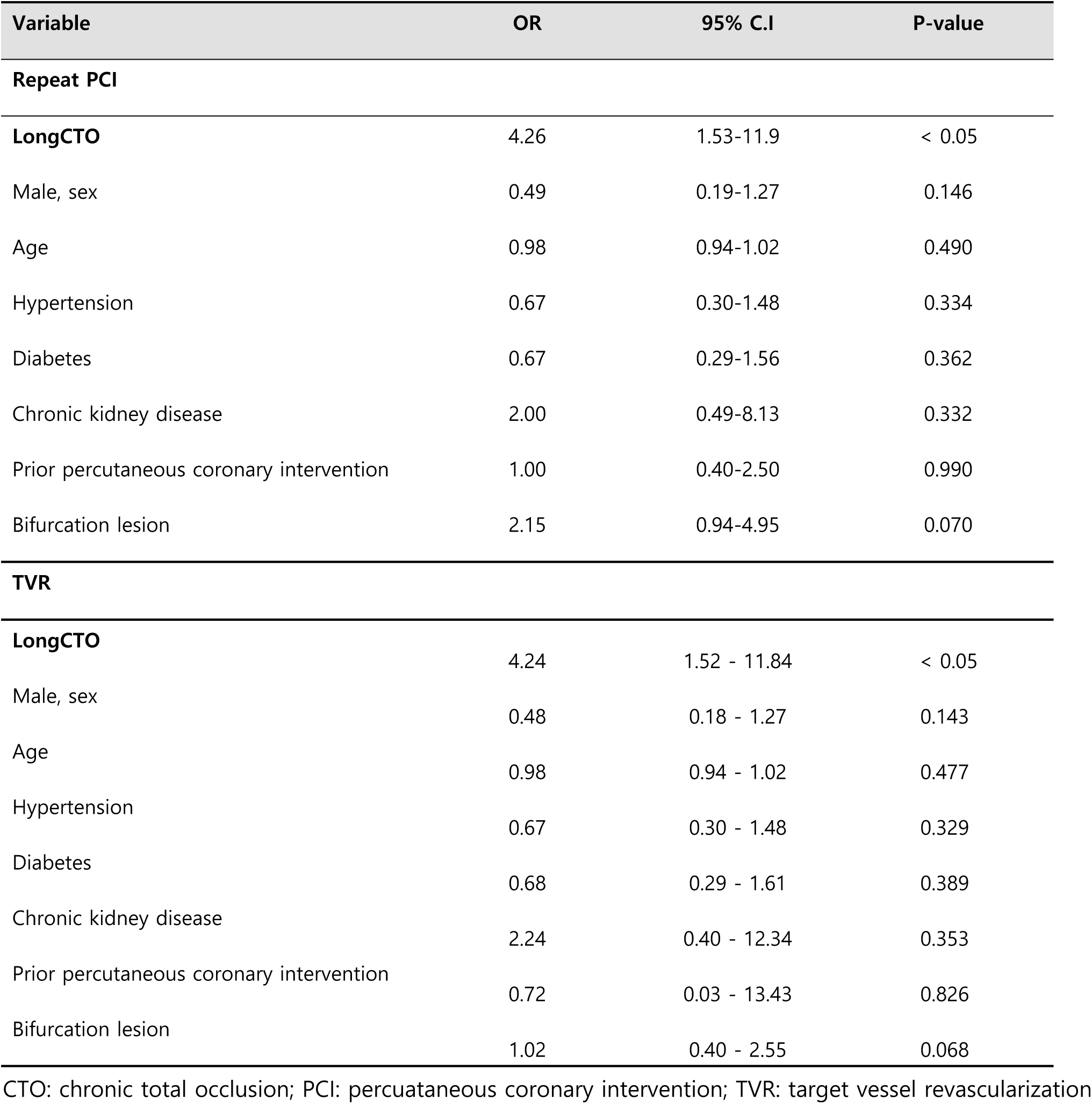
Multivariate Analysis of Risk Factor for Repeat PCI and TVR

### Angiographic outcomes

The data of angiographic follow-up at 6-12 months was obtained in 99 of 159 patients (62%) with the long CTO group and 45 of 76 patients (59%) with the short CTO group (Table 5).

**Table 5.**
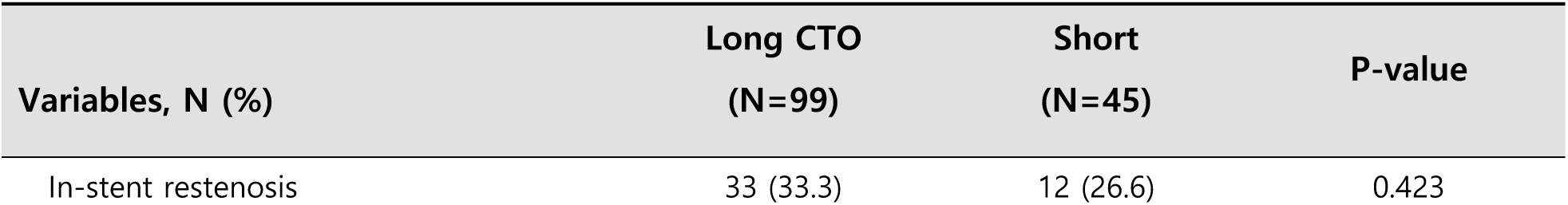
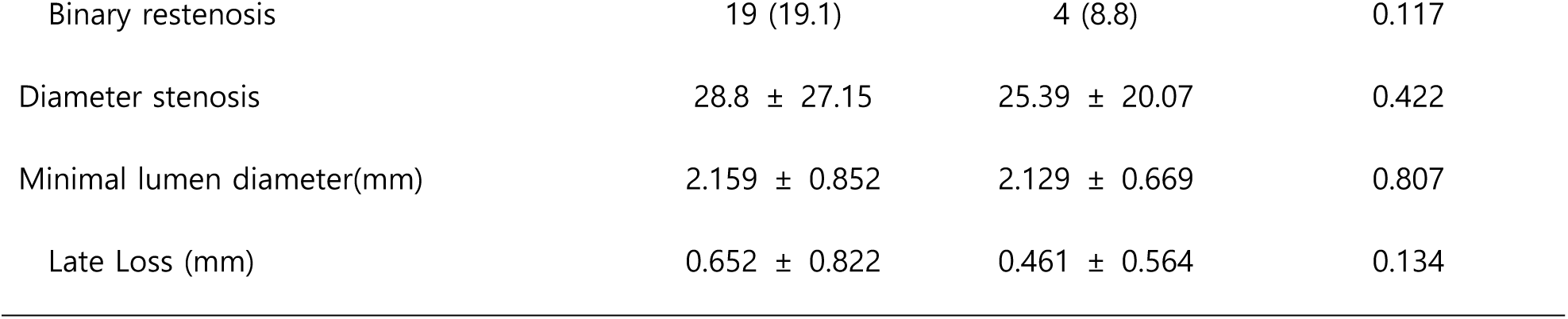
Six-Month Angiographic Outcomes.

Angiographic outcomes were not different between two groups including the incidence of ISR (33.3% vs. 26.6%, p = 0.42), binary restenosis (19.1% cs. 8.8%, p = 0.12) and LL (0.65 ± 0.82 mm vs. 0.46 ± 0.56 mm, p = 0.13).

## Discussion

The main findings of this study were as follows: 1) In the long CTO group, the incidence of prior PCI and history of MI was more frequent compared with the short CTO group. 2) In-hospital complications were similar between the two groups except intimal dissection was more frequent in the long CTO group. 3) Regardless of CTO lesion length, both long and short CTO groups had similar major clinical outcomes including cardiac death and MI and MACE up to 2 years except repeat PCI, predominantly TVR was higher in the long CTO group.4) The long and short CTO groups showed similar angiographic outcomes at 6 months.

In the DES era, successful CTO PCI was not just associated with improved clinical outcomes including cardiac mortality but also a predictor of a reduced rate of coronary artery bypass grafting (CABG) compared with failed CTO PCI[6-9]. Furthermore, with increasing of experience and techniques, procedural success rate of CTO PCI was reported even up to 90%[10, 11]. Nowadays, success rate or technical problems is not a major issue of CTO intervention anymore. However, despite improvement of techniques and success rate of CTO PCI and development of stent technology, clinical and angiographic outcomes of CTO was worse compared with non-CTO disease [12-14]. Also treatment of non-CTO disease, lesion length of the coronary artery disease is a major predictor of MACE and restenosis after stent implantation with DES[15-17].

When we started to investigate the impact of CTO lesion length on clinical outcomes following successful PCI, we carefully predicted and expected that the adverse clinical outcomes including MACE and angiographic adverse outcomes would be more frequent in the long CTO group. The main reasons were; 1) the long CTO group has worse clinical risk factors related to adverse clinical outcomes such as history of prior MI and PCI compared with the short CTO group [18, 19]. 2) Also, use of newer generation DES was dominant in the short CTO group and paclitaxel-eluting stent was frequently used in the long CTO group.

Procedural complications were similar between the two groups except dissection. In the long CTO group, the CTO guide wire passage into false channel would be more frequent than the Short CTO group. However, In-hospital outcomes were not different between the two groups. This result support the procedural safety and efficacy between the two groups during the hospitalization period.

A lot of data support that newer generation DES is associated with superior cardiovascular outcomes following PCI [20, 21]. However, in the present study, the incidence of mortality including cardiac death, MI, and MACE was not different between the two groups. Just the incidences of repeat PCI and TVR were more frequent in the long CTO group. Despite of long CTO was an important independent predictor for repeat PCI (18.8% vs. 7.8%, p < 0.05) and TVR (OR; 4.23, CI 1.52 – 11.8, p < 0.05) in multivariate analysis, overall two-year major clinical outcome results suggest PCI in the long CTO lesion with DES is similarly safe as compared with CTO PCI in the short CTO lesion.

In consideration of our results, the higher incidence of TVR in the long CTO group did not translated into individual critical hard endpoint such as mortality and cardiac death. This different results between hard end points and revascularization incidence in the treatment of CTO was reported previously from the j-Cypher Registry [13]. It is difficult to explain the exact mechanism, however, the presence of less myocardial viability and previously developed collateral channels may play important roles [22, 23].These two factors in the long CTO lesions could lessen the harmful physiologic effect of restenosis in the target vessel.

In the present study, six-month angiographic outcomes between two groups were statistically similar. However, data showed numerically higher trend in the incidence of adverse outcomes such as binary restenosis (19.1% vs. 8.8%, p=0.118). Compared with the results of Hoye A et al’s study, binary restenosis rate (9.1%) of the Short CTO group was similar with our result[24]. In consideration of small patient’s number and follow-up period, we carefully predict the incidence of binary restenosis will be higher in the long CTO group with the time. Also, higher incidence of repeat PCI and TVR during two-year follow-up period support this hypothesis.

Though vessel size was smaller in the short CTO group compared with the long CTO group (3.11 ± 0.43 mm vs. 2.92 ± 0.42, p < 0.05), both groups had enough large vessel diameter on the basis of previous standard categorized as a small vessel diameter[1, 25-27]. Thus this difference of vessel diameter did not affect the angiographic and clinical outcomes significantly.

In this study, there were several study limitations. First, present study is not a randomized study and a single center registry but a data was collected prospectively. And the baseline clinical, lesion and procedural characters were slightly different between the two groups. Also, use of newer generation DES was frequent in the short CTO group and first generation stent was frequently used in the long CTO group. In spite of strict statistical adjustment, there could be unmeasured confounders and analytical bias. Second, follow-up coronary angiography was not routinely performed in all patients. We decided follow-up angiography mainly depends on ischemic symptoms or by physician’s discretion regardless of patient’s symptoms. Well developed collaterals and myocardial viability could influence the ischemic symptoms. So we could not avoid some confounders. Third, the sample size was relatively small, and although we could not find the differences in angiographic outcomes between two groups, the study did not have strong power to prove a difference in angiographic events. Finally, two-year follow-up period was not long enough to evaluate long-term safety and efficacy issues. Despite some limitations, our study have investigated with real world CTO cohort of patients who was collected prospectively. However, long-term study with larger numbers of patients will be required to get final conclusion.

## Conclusions

The safety profile, mid-term angiographic and long-term clinical outcomes were similar between the long CTO and the short CTO group except higher incidence of repeat PCI in the long CTO group despite of DES implantation. Long-term clinical study with larger study population will be necessary to elucidate the final conclusion.

## Notes

– Conflicts of Interest: The authors have no financial conflicts of interest.

## References

1. Habara S, Mitsudo K, Goto T, Kadota K, Fujii S, Yamamoto H, et al. The impact of lesion length and vessel size on outcomes after sirolimus-eluting stent implantation for in-stent restenosis. Heart. 2008;94(9):1162–5.

2. Kastrati A, Dibra A, Mehilli J, Mayer S, Pinieck S, Pache J, et al. Predictive factors of restenosis after coronary implantation of sirolimus- or paclitaxel-eluting stents. Circulation. 2006;113(19):2293–300.

3. Serruys PW, Silber S, Garg S, van Geuns RJ, Richardt G, Buszman PE, et al. Comparison of zotarolimus-eluting and everolimus-eluting coronary stents. N Engl J Med. 2010;363(2):136–46.

4. Li R, Yang S, Tang L, Yang Y, Chen H, Guan S, et al. Meta-analysis of the effect of percutaneous coronary intervention on chronic total coronary occlusions. J Cardiothorac Surg. 2014;9:41.

5. Stone GW, Kandzari DE, Mehran R, Colombo A, Schwartz RS, Bailey S, et al. Percutaneous recanalization of chronically occluded coronary arteries: a consensus document: part I. Circulation. 2005;112(15):2364–72.

6. Mehran R, Claessen BE, Godino C, Dangas GD, Obunai K, Kanwal S, et al. Long-term outcome of percutaneous coronary intervention for chronic total occlusions. JACC Cardiovasc Interv. 2011;4(9):952–61.

7. de Labriolle A, Bonello L, Roy P, Lemesle G, Steinberg DH, Xue Z, et al. Comparison of safety, efficacy, and outcome of successful versus unsuccessful percutaneous coronary intervention in “true” chronic total occlusions. Am J Cardiol. 2008;102(9):1175–81.

8. Prasad A, Rihal CS, Lennon RJ, Wiste HJ, Singh M, Holmes DR, Jr. Trends in outcomes after percutaneous coronary intervention for chronic total occlusions: a 25-year experience from the Mayo Clinic. J Am Coll Cardiol. 2007;49(15):1611–8.

9. Suero JA, Marso SP, Jones PG, Laster SB, Huber KC, Giorgi LV, et al. Procedural outcomes and long-term survival among patients undergoing percutaneous coronary intervention of a chronic total occlusion in native coronary arteries: a 20-year experience. J Am Coll Cardiol. 2001;38(2):409–14.

10. Morino Y, Kimura T, Hayashi Y, Muramatsu T, Ochiai M, Noguchi Y, et al. In-hospital outcomes of contemporary percutaneous coronary intervention in patients with chronic total occlusion insights from the J-CTO Registry (Multicenter CTO Registry in Japan). JACC Cardiovasc Interv. 2010;3(2):143–51.

11. Werner GS, Hochadel M, Zeymer U, Kerber S, Schumacher B, Grube E, et al. Contemporary success and complication rates of percutaneous coronary intervention for chronic total coronary occlusions: results from the ALKK quality control registry of 2006. EuroIntervention. 2010;6(3):361–6.

12. Dangas GD, Claessen BE, Caixeta A, Sanidas EA, Mintz GS, Mehran R. In-stent restenosis in the drug-eluting stent era. J Am Coll Cardiol. 2010;56(23):1897–907.

13. Kato M, Kimura T, Morimoto T, Nishikawa H, Uchida F, Suzuki H, et al. Comparison of five-year outcome of sirolimus-eluting stent implantation for chronic total occlusions versus for non-chronic total occlusion (from the j-Cypher registry). Am J Cardiol. 2012;110(9):1282–9.

14. Valenti R, Vergara R, Migliorini A, Parodi G, Carrabba N, Cerisano G, et al. Predictors of reocclusion after successful drug-eluting stent-supported percutaneous coronary intervention of chronic total occlusion. J Am Coll Cardiol. 2013;61(5):545–50.

15. Claessen BE, Smits PC, Kereiakes DJ, Parise H, Fahy M, Kedhi E, et al. Impact of lesion length and vessel size on clinical outcomes after percutaneous coronary intervention with everolimus-versus paclitaxel-eluting stents pooled analysis from the SPIRIT (Clinical Evaluation of the XIENCE V Everolimus Eluting Coronary Stent System) and COMPARE (Second-generation everolimus-eluting and paclitaxel-eluting stents in real-life practice) Randomized Trials. JACC Cardiovasc Interv. 2011;4(11):1209–15.

16. Quadri G, D’Ascenzo F, Bollati M, Moretti C, Omede P, Sciuto F, et al. Diffuse coronary disease: short- and long-term outcome after percutaneous coronary intervention. Acta Cardiol. 2013;68(2):151–60.

17. Rathore S, Terashima M, Katoh O, Matsuo H, Tanaka N, Kinoshita Y, et al. Predictors of angiographic restenosis after drug eluting stents in the coronary arteries: contemporary practice in real world patients. EuroIntervention. 2009;5(3):349–54.

18. Motivala AA, Tamhane U, Ramanath VS, Saab F, Montgomery DG, Fang J, et al. A prior myocardial infarction: how does it affect management and outcomes in recurrent acute coronary syndromes? Clin Cardiol. 2008;31(12):590–6.

19. Stolker JM, Cohen DJ, Kennedy KF, Pencina MJ, Lindsey JB, Mauri L, et al. Repeat revascularization after contemporary percutaneous coronary intervention: an evaluation of staged, target lesion, and other unplanned revascularization procedures during the first year. Circ Cardiovasc Interv. 2012;5(6):772–82.

20. Kaul U, Bangalore S, Seth A, Arambam P, Abhaichand RK, Patel TM, et al. Paclitaxel-Eluting versus Everolimus-Eluting Coronary Stents in Diabetes. N Engl J Med. 2015;373(18):1709–19.

21. Kedhi E, Joesoef KS, McFadden E, Wassing J, van Mieghem C, Goedhart D, et al. Second-generation everolimus-eluting and paclitaxel-eluting stents in real-life practice (COMPARE): a randomised trial. Lancet. 2010;375(9710):201–9.

22. Werner GS. The role of coronary collaterals in chronic total occlusions. Curr Cardiol Rev. 2014;10(1):57–64.

23. Werner GS, Emig U, Mutschke O, Schwarz G, Bahrmann P, Figulla HR. Regression of collateral function after recanalization of chronic total coronary occlusions: a serial assessment by intracoronary pressure and Doppler recordings. Circulation. 2003;108(23):2877–82.

24. Hoye A, Tanabe K, Lemos PA, Aoki J, Saia F, Arampatzis C, et al. Significant reduction in restenosis after the use of sirolimus-eluting stents in the treatment of chronic total occlusions. J Am Coll Cardiol. 2004;43(11):1954–8.

25. Elezi S, Dibra A, Mehilli J, Pache J, Wessely R, Schomig A, et al. Vessel size and outcome after coronary drug-eluting stent placement: results from a large cohort of patients treated with sirolimus- or paclitaxel-eluting stents. J Am Coll Cardiol. 2006;48(7):1304–9.

26. Mauri L, Orav EJ, Kuntz RE. Late loss in lumen diameter and binary restenosis for drug-eluting stent comparison. Circulation. 2005;111(25):3435–42.

27. van der Heijden LC, Kok MM, Danse PW, Schramm AR, Hartmann M, Lowik MM, et al. Small-vessel treatment with contemporary newer-generation drug-eluting coronary stents in all-comers: Insights from 2-year DUTCH PEERS (TWENTE II) randomized trial. Am Heart J. 2016;176:28–35.

